# Prorenin from renal tubules is a major driver of diabetic kidney disease

**DOI:** 10.1101/2022.09.26.509473

**Authors:** Anna Federlein, Claudia Lehrmann, Philipp Tauber, Vladimir Todorov, Frank Schweda

## Abstract

The renin-angiotensin-system (RAS) plays a critical role in diabetic nephropathy, and inhibitors of the RAS are central components in its therapy. Renin circulates in the blood in its active form but also in the form of its enzymatically inactive precursor prorenin. In humans, the plasma prorenin concentration exceeds that of active renin several times. While the plasma concentration of active renin is unchanged or even reduced in diabetic patients, the prorenin concentration increases significantly and it has been shown that high prorenin levels are associated with diabetic microvascular damage. Why prorenin increases in diabetes while active renin is reduced is unclear. Previous studies suggest that there may be formation of prorenin in the tubular system of diabetic kidneys. To investigate the functional consequences of possible tubular renin formation in diabetes, we generated mice with inducible tubule-specific deletion of the renin gene (tubule-renin KO).

Under control conditions, tubule-renin KO had no apparent phenotype and plasma and tissue levels of renin and prorenin were similar to those of mice with intact tubular renin (control mice). In control mice type-1 diabetes (streptozotocin, STZ) for 8 to 12 weeks stimulated tubular renin mRNA and protein expression, especially in distal nephron segments including collecting ducts. This increase in renin synthesis in renal tubules was markedly attenuated or even absent in tubule-renin KO mice. Similar to diabetic patients, plasma prorenin was markedly elevated in diabetic control mice. This stimulation of plasma prorenin was absent in mice with tubule-specific deletion of the renin gene. Moreover, the high prorenin levels in renal tissue, which were observed in diabetic control mice, were markedly reduced in tubule-renin KO. Noteworthy, plasma renin activity was not reduced in tubule-renin KO compared with controls, suggesting that renal tubules of diabetic mice mainly release prorenin. Kidneys of control mice with intact tubular renin showed classical signs of diabetic renal damage, such as albuminuria, mesangial expansion, fibrosis, inflammation and capillary rarefaction. All of these parameters were significantly ameliorated in tubule-renin KO, indicating that tubular renin, most likely in its prorenin form, significantly aggravates renal damage in diabetes.

These data provide clear evidence for the first time that the tubular renin system is an additional source for circulating prorenin in diabetes, hereby providing an explanation for the paradox regulation of active renin and prorenin in diabetes. Moreover, the data show that tubular renin markedly contributes to the progression of diabetic nephropathy.

## Introduction

The renin-angiotensin-system not only plays a central role in the regulation of salt and water homeostasis and the blood pressure but also in cardiovascular and renal diseases. Accordingly, RAS blockers, such as ACE-inhibitors or AT1-receptor antagonists, are first line therapeutics in these diseases including diabetic nephropathy.

Renin is synthesized in and released from renin producing juxtaglomerular (JG cells) that are located in the vessel walls of afferent arterioles. Main factors stimulating renin release from JG cells are renal perfusion pressure, salt deficiency and high sympathetic nerve activity (Schweda, 2015). In addition to JG cells, tubular cells express renin. While renin expression is very low to undetectable under normal conditions, it is markedly stimulated in renal collecting ducts in ANGII-dependent forms of arterial hypertension (Prieto-Carrasquero et al., 2008; Gonzalez et al., 2011). Furthermore, mice with a genetically induced salt-losing nephropathy show a drastic stimulation of the circulating RAS together with clear tubular renin staining (Grill et al., 2016). Since angiotensinogen can enter the tubular lumen by glomerular filtration or by secretion from proximal tubular cells and the angiotensin converting enzymes (ACE 1 and ACE 2) are expressed in the tubular system, the nephron contains all necessary components to form a local RAS (Pohl et al., 2010). The concentrations of ANGII in the tubular fluid and in the renal tissue both markedly exceed plasma levels, suggesting the existence of an intrarenal RAS with local ANGII formation (Seikaly et al., 1990; van Kats et al., 2001). Since ANGII can directly stimulate Na^+^-reabsorption via activation of the epithelial Na^+^-channel ENaC, locally produced ANGII could be involved in blood pressure regulation via elevating the extracellular volume (Mamenko et al., 2012). Since tubular renin expression is stimulated in ANGII-dependent forms of arterial hypertension, this mechanism should gain relevance in these conditions. In fact, ANGII-dependent arterial hypertension is attenuated in mice with collecting duct specific deletion of renin (Ramkumar et al., 2014) while this is not the case in DOCA-salt hypertension (Song et al., 2016).

Renin is synthesized in JG cells in its inactive pro-form prorenin. Prorenin can either be released continuously or be packed into secretory vesicles in which it is processed into active renin via cleavage of the prosequence. While the release of active renin from the secretory granules is rapidly regulated, the constitutive pathway of prorenin release is determined primarily by the prorenin synthesis rate (Schweda and Kurtz, 2011). Prorenin can bind to the (pro)renin receptor (PRR) (Nguyen et al., 2002), which is a multifunctional protein. Within the kidney, PRR is predominantly expressed in the collecting duct (Advani et al., 2009; Daryadel et al., 2016). Upon binding to the receptor, prorenin gets enzymatically active and can contribute to the local production of ANGII, at least *in vitro* (Nguyen et al., 2002). Next to this role as a receptor, the PRR protein is an accessory subunit of the vacuolar H^+^-ATPase (Advani et al., 2009). The *in vivo* relevance of the PRR as a component of the RAS is highly debated on the background of conflicting results from studies using genetic models or pharmacological blockers of the PRR (Kaneshiro et al., 2007; Feldt et al., 2008; Ramkumar et al., 2016; Trepiccione et al., 2016). Therefore, to date the functional relevance of prorenin binding to the PRR is not completely resolved.

As mentioned above, RAS inhibitors are a cornerstone therapy for diabetic nephropathy. On the background of the therapeutic effects of RAS inhibitors, it is counterintuitive that plasma renin activity is low to normal in diabetic patients (Fernandez-Cruz et al., 1981; Price et al., 1999; Chiarelli et al., 2001). Intriguingly, plasma prorenin levels are elevated in diabetic patients and high prorenin concentrations are associated with microvascular complications such as retinopathy or nephropathy and even precede and predict microalbuminuria (Luetscher et al., 1985; Wilson and Luetscher, 1990; Chiarelli et al., 2001). Studies in rats suggested that elevated prorenin in diabetes might derive from renal tubules (Kang et al., 2008). Based on these studies an activation of the intrarenal RAS in diabetes has been suggested. In line, urinary angiotensinogen and renin levels are elevated in diabetes (van den Heuvel et al., 2011; Park et al., 2015). However, data from a recent study rather indicated glomerular filtration and altered proximal reabsorption of renin as the underlying mechanism of elevated urinary renin excretion (Tang et al., 2019). In order to directly address the question whether renal tubules are a source of renin and/or prorenin in diabetes and to investigate the functional role of tubular renin, we generated mice with tubule-specific deletion of renin and subjected them to streptozotocin diabetes.

## Methods

### Generation of mice with floxed renin allele

Mutant mouse embryonic stem cells with targeted Ren1c locus were purchased from the European Mouse Mutant Cell Repository - EuMMCR (Helmholtz Zentrum München, Germany). The targeting vector used for homologous recombination of ES cells contained 5’ and 3’ homology arms as well as a central targeting cassette with a neo-resistance gene and a lacZ reporter, flanked by FRT flippase recombination sites followed by two loxP recombination sites flanking exons 5-8 of the mouse *Ren1c* gene. Ren1c floxed mice were generated in the Transgenic Core Facility, Max Planck Institute of Molecular Cell Biology and Genetics (MPI-CBG), Dresden. JM8A1.N3 cells were injected into 8-cell stage C57Bl/6NCrl embryos by Laser assisted injection (XY-Clone Hamilton Thorne). Three different clones were cultured and injected; one Clone (D03) gave high chimeras and germline transmission. Chimeras were crossed to C57Bl/6NCrl mice and were screened by PCR using primers:

Ren1-5’arm: 5’-CAG CCT CCT TGG CAG CTT CTA GCC-3’
Ren1-3’arm: 5’-ACT GTC AAC ACC TCT ATG CTT GGG-3’
LAR3: 5’-CAA CGG GTT CTT CTG TTA GTC C-3’

The PCR-products sizes are 519 bp for the wildtype allele and 392 bp for the targeted allele.

Heterozygous mice carrying the targeting construct were crossed to flpo (improved flippase) deleter strain to excise the targeting cassette (Kranz et al., 2010). Flpo recombination was screened by PCR using primers Ren1-5’arm and Ren1-3’arm. The PCR-products sizes are 519 bp for the wildtype allele and 667 bp for the floxed allele.

After global deletion of the targeting cassette the flpo transgene was outbred. The resulting line (Ren1^flox^) has two loxP sites that flank exons 5-8 of the mouse *Ren1c* gene. Upon cre-mediated recombination the floxed exons are excised resulting in loss of the second lobe of the renin molecule active site and consecutive loss of enzymatic activity.

### Generation of inducible tubule specific reporter mice

We used the inducible Pax8-TetO-Cre mouse line that directs expression of the reverse tetracycline-dependent transactivator to all tubular segments ((B6.Cg-Tg(Pax8-rtTA2S*M2)1Koes/J8) (Traykova-Brauch et al., 2008). These mice were crossed with double-fluorescent Cre reporter mice (Rosa mTmG (Gt(ROSA)26Sortm4(ACTB-tdTomato,-EGFP)Luo/J)) (Muzumdar et al., 2007). The resulting mice express membrane-targeted tomato (mT, red fluorescence) in all cells prior to Cre-mediated recombination. After induction of the transactivator and activation of Cre-recombinase, cells with Cre-activity express the membrane-targeted green fluorescent protein (mG, green fluorescence), while cells without Cre-activity remain red.

### Generation of an inducible tubule-specific renin KO mouse

Male mice at the age of 12 to 16 weeks were included into the experiments. Mice with a tubule-specific deletion of renin (“tubule-renin KO”) were generated by crossing the inducible Pax8-TetO-Cre mice (see above) with the newly generated Ren1^flox^ mice mentioned above. Pax8-TetO-Cre x Ren1^WT/WT^ mice and Ren1^flox/flox^ mice (without Pax8-Cre) were considered as controls, Pax8-TetO-Cre x Ren1^flox/flox^ as tubule-specific knockouts. Cre-mediated recombination was induced by doxycycline (2 mg/ml via drinking water for 21 days, PanReac Applichem, Darmstadt, Germany). Also control mice were treated with doxycycline. To mask the bitter taste of doxycycline, 5% sucrose were added to the water. A 2week washout period was granted before the start of experiments.

All mice were bred and housed under specific pathogen-free conditions and food and water were provided ad libitum. All animal experiments were conducted according to the German law for animal care and approved by the local authorities (54-2532.1-20/14 & 55.2-2532.2-1107-14).

### Streptozotocin diabetes mellitus model

Hyperglycemia was induced by injections of streptozotocin (50 mg/kg b.w., i.p., for 5 consecutive days). STZ was dissolved in a 50 mM sodium citrate buffer (pH 4.5) immediately prior to the injection. Mice were fasted for at least four hours prior to the injections. Glucose levels were determined in a blood drop taken from the facial vein 7 days after STZ-treatment using a glucometer (Contour, Bayer, Basel, Switzerland). Mice with blood glucose level higher than 15 mmol / l were classified as diabetic.

### Determination of Plasma Renin Concentration (PRC)

Blood samples were taken by puncture of the facial vein and collected in haematocrit-capillaries. The quantitative determination of renin activity is based on the generation of angiotensin I in the presence of excess angiotensinogen as described previously (Tauber et al., 2021). The generated angiotensin I [ng/ml x h^-1^] was determined by enzyme immunoassay (IBL International, Hamburg, Germany).

### Determination of Renal Renin Content

Renal renin content was determined by measuring the capacity of homogenized kidney tissue to generate angiotensin I in the presence of excess renin substrate. Kidneys were dissected on ice into cortex, outer medulla and papilla and snap frozen in liquid nitrogen. Frozen tissue samples were weighted and homogenized in a buffer containing protease inhibitors (5% (vol/vol) glycerol, 0.1 mM phenylmethylsulfonyl fluoride, 10 mM EDTA, and 0.1 mM 4-(2-aminomethyl) benzenesulfonyl fluoride) for 30 s (Ultra-Turrax, IKA Labortechnik) and centrifuged at 4°C at 14,000 *g* for 5 min. Supernatants were incubated with saturating concentrations of rat renin substrate, and the generated angiotensin I was determined by EIA as mentioned above.

### Determination of Urinary Renin Excretion

Spot urine was collected at baseline conditions and 12 weeks after the induction of STZ diabetes and the angiotensin I-to-creatinine ratio was determined. Urine samples were incubated with rat renin substrate for 1.5 hours at 37°C and the generated angiotensin I was detected by EIA (IBL International, Hamburg, Germany). Creatinine was measured by a colorimetric assay (BioAssay Systems, Hayward, CA).

### Determination of plasma, urine and tissue prorenin

The prorenin concentration was calculated as the difference between the total and active renin. To determine the total renin concentration, prorenin was activated to renin by trypsinization (Schmid et al., 2013). For each sample type (plasma, urine, renal tissue) the optimal trypsin concentration and duration of incubation, resulting in highest renin activity levels, had been identified before. The reaction was stopped with soybean trypsin inhibitor. The samples were incubated with rat renin substrate and the generation of angiotensin I was measured using an enzyme immunoassay (IBL International, Hamburg, Germany).

### Glomerular filtration rate

The glomerular filtration rate (GFR) was measured in conscious mice by using the transdermal GFR technology as described previously (MediBeacon Inc., Mannheim, Germany) (Schreiber et al., 2012; Tauber et al., 2021).

### Blood pressure measurement by the tail-cuff method

Mice were conditioned by placing them in the holding devices for 7 consecutive days before the first measurement was performed. Subsequently, eight blood pressure and heart rate values per mouse were determined per day and averaged for a total of 5 consecutive days.

### Urinary Albumin/creatinine ratio

Spot urine was collected at baseline conditions and 12 weeks after the induction of STZ diabetes. Albumin was quantified with a specific mouse albumin EIA according to the manufacturer’s instructions (Dunn Labortechnik, Asbach, Germany). A colorimetric assay (BioAssay Systems, Hayward, CA) was used for the determination of creatinine in the urine.

### Histologic Analysis of Kidney Sections

Kidneys were perfusion-fixed with 4% paraformaldehyde, halved and stored in 70% methanol overnight. Subsequently, the fixed kidneys were dehydrated, embedded in paraffin and sectioned in 5-μm-thick sections. The kidney sections were either stained with periodic acid-Schiff (PAS) reaction using a standard protocol or with Masson Trichrome according to the manufacturer’s protocol (Sigma, Taufkirchen, Germany) and examined by light microscopy (Axio Observer 7, Carl Zeiss, Germany).

### PAS

Glomerular and mesangial areas of at least 50 glomeruli per mouse were determined in PAS-stained sections using ZEN lite software by two independent investigators, who were blinded regarding genotype and treatment.

### Masson Goldner

Stained areas per kidney section were quantified in an unbiased manner with Zeiss ZEN Intellesis Image Segmentation software as described previously (Tauber et al., 2021).

### Immunofluorescence Staining

Sections were blocked with 10% horse serum / 1% bovine serum albumin in phosphate-buffered saline and incubated at 4°C overnight with the following primary antibodies: polyclonal rabbit anti-renin (dilution 1:800) (OACA02177, Avia Systems Biology, San Diego, USA), polyclonal goat anti-aquaporin-2 (dilution 1:200) (sc-9882, Santa Cruz Biotechnology, Santa Cruz, CA) monoclonal mouse anti-calbindin (dilution 1:500) (D-28k, Swant, Marly, Switzerland), monoclonal anti-mouse α-SMA antibody (dilution 1:600) (ab7817, Abcam plc, Cambridge, United Kingdom), polyclonal goat anti-albumin (dilution 1:500) (ab 19194, Abcam, Cabridge, UK), polyclonal chicken anti-GFP (dilution 1:600) (ab13970, Abcam, Cambridge, UK), polyclonal goat anti-CD31 antibody (dilution 1:200) (AF3628, R&D Systems, USA), monoclonal rat anti-F4/80 antibody (dilution 1:200) (clone Cl:A3-1, MCA497G, Biorad Laboratories, Germany) After multiple washes, the appropriate secondary antibody was incubated at room temperature for 90 minutes: Alexa Flour 488-conjugated AffiniPure donkey anti-rabbit IgG (711-545-152; Jackson ImmunoResearch), DyLight 488-conjugated AffiniPure donkey anti-chicken IgG (703-225-155; Jackson ImmunoResearch), DyLight 488-conjugated AffiniPure donkey anti-goat IgG (705-485-147; Jackson ImmunoResearch), Rhodamine (TRICT)-conjugated AffiniPure donkey anti-goat IgG (705-025-147), Rhodamine (TRICT)-conjugated AffiniPure donkey anti-mouse IgG (715-025-150; Jackson ImmunoResearch), Rhodamine (TRICT)-conjugated AffiniPure donkey anti-rat IgG (712-025-153; Jackson ImmunoResearch). For nuclear staining, 4’, 6-Diamidino-2-phenylindole dihydrochloride (dilution 1:400) (sc-3598 1μg/ml) was applied for 90 minutes together with the secondary antibody. Slides were mounted with Dako Glycergel (C0563, Dako, Glostrup, Denmark) and images were collected using an Axio Observer 7 fluorescence microscope and a confocal microscope (Zeiss LSM 710).

Cryosections were used for the observation of autofluorescence and costaining with renin in kidneys of mG/mT reporter mice. For this, the kidneys were perfusion-fixed as described above and stored in 1% paraformaldehyde / 18% sucrose in phosphate-buffered saline overnight. After a brief immersion in −40°C cold methyl butane, the kidneys were snap frozen in liquid nitrogen. 5μm-thick-sections were washed with phosphate-buffered saline, permeabilized with 0,1% sodium dodecyl sulfate in phosphate-buffered saline and blocked with 5% bovine serum albumin / 0,04% Triton-X in phosphate-buffered saline. After incubation with the primary polyclonal rabbit anti-renin antibody (dilution 1:800) (OACA02177, Avia Systems Biology, San Diego, USA) at 4°C overnight, the sections were washed and incubated for 60 min with the secondary antibody Cy5-conjugated AffiniPure donkey anti-rabbit IgG (711-175-152; Jackson ImmunoResearch). For nuclear staining, 4’, 6-Diamidino-2-phenylindole dihydrochloride was added (dilution 1:400) (sc-3598 1μg/ml). The slides were mounted with Dako Glycergel (Dako, Glostrup, Denmark).

### Quantification of F4/80+ macrophages

The number of intrarenal macrophages was determined by F4/80 fluorescent staining. F4/80 positive stained cells were counted on at least 25 pictures taken with an Axio Observer 7 fluorescence microscope (original magnification 200x) of each kidney. Cell counting was performed in a blinded manner using ImageJ (National Institutes of Health, version 1.52a)

### Capillary quantification - CD31 staining

To analyze the capillary density the endothelial cell marker CD 31 was used. Pictures of 30 visual fields of each kidney were taken with an Axio Observer 7 fluorescence microscope (original magnification 400x) and the number of peritubular capillaries (PTCs) and tubules was counted. The data shown in the results are presented as the ratio of the number of PTCs to tubuli. Counting was performed by 2 persons independently in a blinded fashion using ImageJ software.

### In situ hybridization via RNAscope

Localization of renin mRNA expression was analyzed using a chromogenic RNAscope Brown 2.5 HD reagent kit and a Renin target probe (Mm-Ren1, Cat#433461 Advanced Cell Diagnostics ACD, Newark, CA), according to the manufacturer’s instructions (Wang and Flanagan et al. 2012). For hybridization, kidneys were perfusion-fixed and further processed as described above. Slices were mounted with VectaMount mounting media (Vector Laboratories, Burlingame, CA) and viewed with an Axio Observer 7 microscope (Carl Zeiss, Germany).

### mRNA Expression by Real-Time PCR

Total RNA was isolated from tissue of the inner medulla, microdissected tubules of the outer medulla and cortex and glomeruli using TRIzol reagent (Biocat GmbH, Heidelberg, Germany). RNA content was quantified by a NanoDrop spectrophotometer. After reverse transcription (Oligo(dT)20 primers and m-MLV RT, Life Technologies, Carlsbad, CA), quantitative real-time PCR was performed using a LightCycler Instument (Roche Diagnostics Corporation) using the following primers: renin forward primer 5’-AGGGGGTGCTAAAGGAGGAA-3’, reverse Primer 5’-GATAATGCTGCGGGTCGCTA-3’;18s: forward primer 5’-AAACGGCTACCACATCCAAG-3’, reverse primer 5’-CCTCCAATGGATCCTCGTTA-3’). Amplification products were analyzed by melting curve analysis and the length of the products were controlled by gel electrophoresis. Data were normalized to 18s or RPL32 as housekeeping genes.

### Human Kidney Biopsies

The human kidney tissue used in this study was obtained from 4 type-1 and 4 type-2 diabetic patients. It was formalin fixed (4% buffered formalin) and paraffin embedded (FFPE), and specimens were provided by the Department of Nephropathology (Prof. Dr. K Amann, Prof. Dr. C. Daniel; Friedrich–Alexander-University, Erlangen–Nuremberg, Germany). The analysis of archived renal tissue was approved by the local Ethics committee.

### Statistics

All data are presented as mean ± SEM. For single-group comparisons, an unpaired student’s t test was used to calculate the level of significance. To analyze the difference between the groups at different time points a two-way ANOVA followed by a Bonferroni post hoc test was used. GraphPad Prism 6.0 (GraphPad Software Inc, CA, US) was used for statistical analysis. P values less than 0.05 were considered statistically significant.

## Results

### Diabetes mellitus induces (pro)renin expression in the tubular system

The antibody used in this study does detect both prorenin and the active renin protein. Therefore, immunostaining does not allow to discriminate between these two forms, and the term (pro)renin, used in the following, does include both forms. In non-diabetic mice (pro)renin protein expression was restricted to its classical position in juxtaglomerular cells (JG cells) and no tubular (pro)renin staining was detectable (Fig. 1A, control). While (pro)renin expression in JG cells was unaltered in diabetic compared with non-diabetic mice, additional clear (pro)renin staining occurred in renal tubules of diabetic mice (Fig.1A, STZ diabetes). Co-staining with segment specific antibodies revealed that (pro)renin was mainly present in collecting ducts and distal tubules (Fig.1A), while other tubular segments did not display robust and reproducible staining (not shown). Tubular (pro)renin expression was not only detectable in diabetic mice but was also in type 1 and type 2 diabetic patients (Fig.1B).

**Figure 1:**
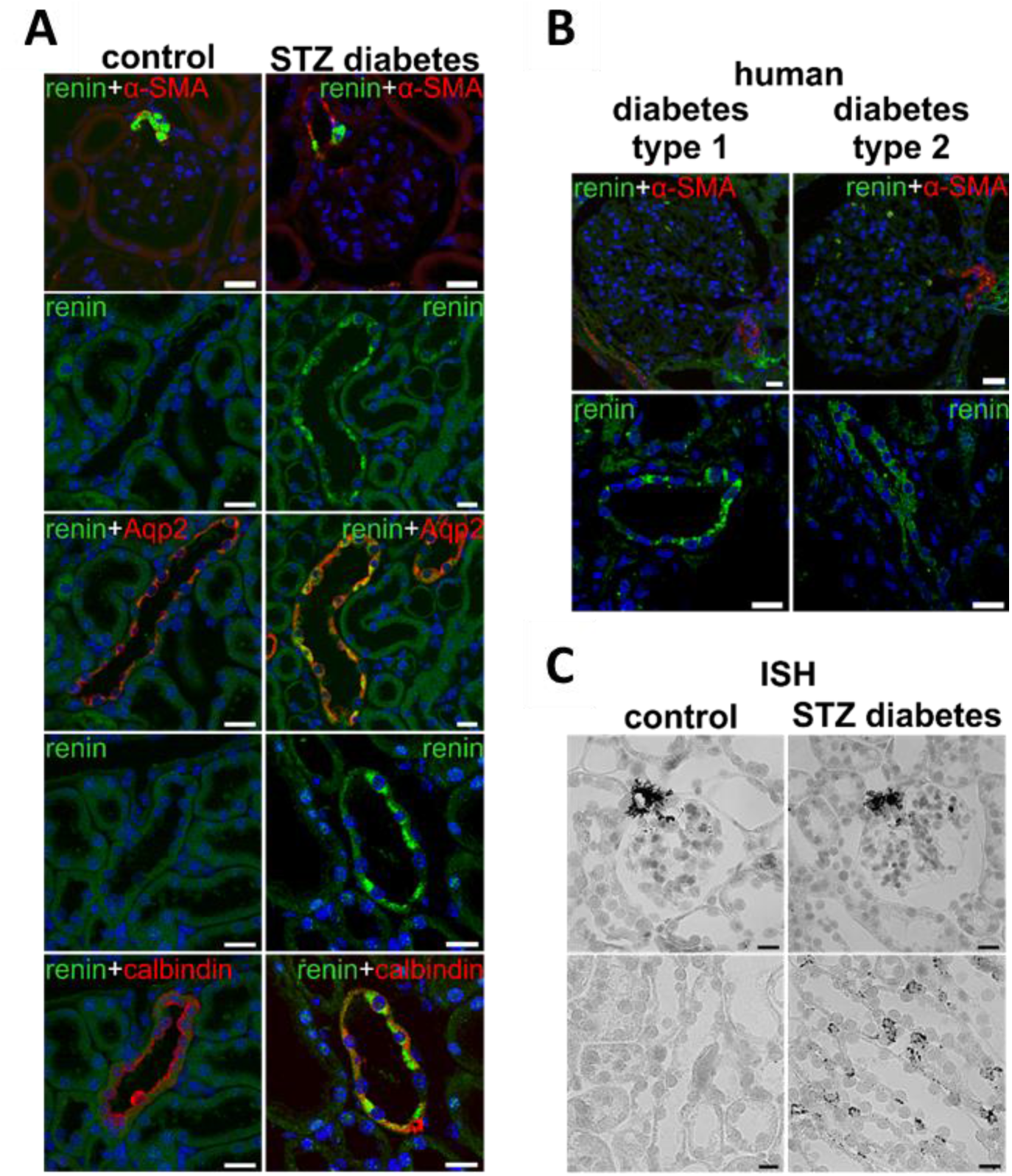
(Pro)renin expression in murine and human kidneys. (A) (Pro)renin staining in kidneys from control mice or mice with STZ-diabetes. Co-staining with αSMA (vascular smooth muscle cells), aquaporin-2 (Aqp2, collecting ducts) and calbindin (distal tubule). (B) (Pro)renin staining in kidneys from type 1 or type 2 diabetic patients: In addition to (pro)renin staining in juxtaglomerular cells (upper panel), tubules were (pro)renin positive. (C) In situ hybridization (RNAscope) in kidneys from control mice or mice with STZ-diabetes. Scale bars: 20 μm.

(Pro)renin protein expression detected by immunofluorescence might result from synthesis in tubular cells or uptake from the tubular or interstitial fluid. In order to test for local production of (pro)renin, preprorenin mRNA expression was determined by RNAscope technology. As expected, marked expression of (pro)renin mRNA was visible in JG cells (Fig. 1C). Moreover, clear signals were observed in tubules of STZ-diabetic but not in non-diabetic mice, clearly supporting the conclusion that a tubular synthesis of (pro)renin exists under diabetic conditions (Fig. 1C).

### Generation of tubule-specific renin knockout mice

In order to test for the functional consequences of the diabetes-induced upregulation of tubular (pro)renin synthesis, mice with specific deletion of renin in renal tubules were generated. Since (pro)renin expression was not restricted to a single tubular segment, deletion of renin expression in the entire tubular system was performed. In order to prevent possible kidney development defects due to the lack of renin, the inducible Pax8-TetO-Cre mouse deleter strain was chosen. Crossing of Pax8-TetO-Cre with mT/mG reporter mice (Muzumdar et al., 2007) and induction with doxycycline resulted in Cre-mediated recombination, identifiable by green GFP fluorescence, in the entire tubular system but neither in the renal vasculature nor, most importantly, in renin producing JG cells (Figure 2A, upper panels). No recombination occurred without induction with doxycycline, so that all cells displayed red fluorescence (Figure 2A, lower panels). Crossing of Pax8-TetO-Cre mice with mice carrying floxed alleles of the renin gene (Pax8-TetO-Cre x Ren ^flox/flox^, termed “tubule-renin KO” throughout this paper) and activation of the transgene by doxycycline resulted in a marked reduction of renin expression in microdissected collecting ducts to 15% of wildtype levels (Figure 2B), while renin expression in glomeruli remained unaltered (Figure 2C).

**Figure 2:**
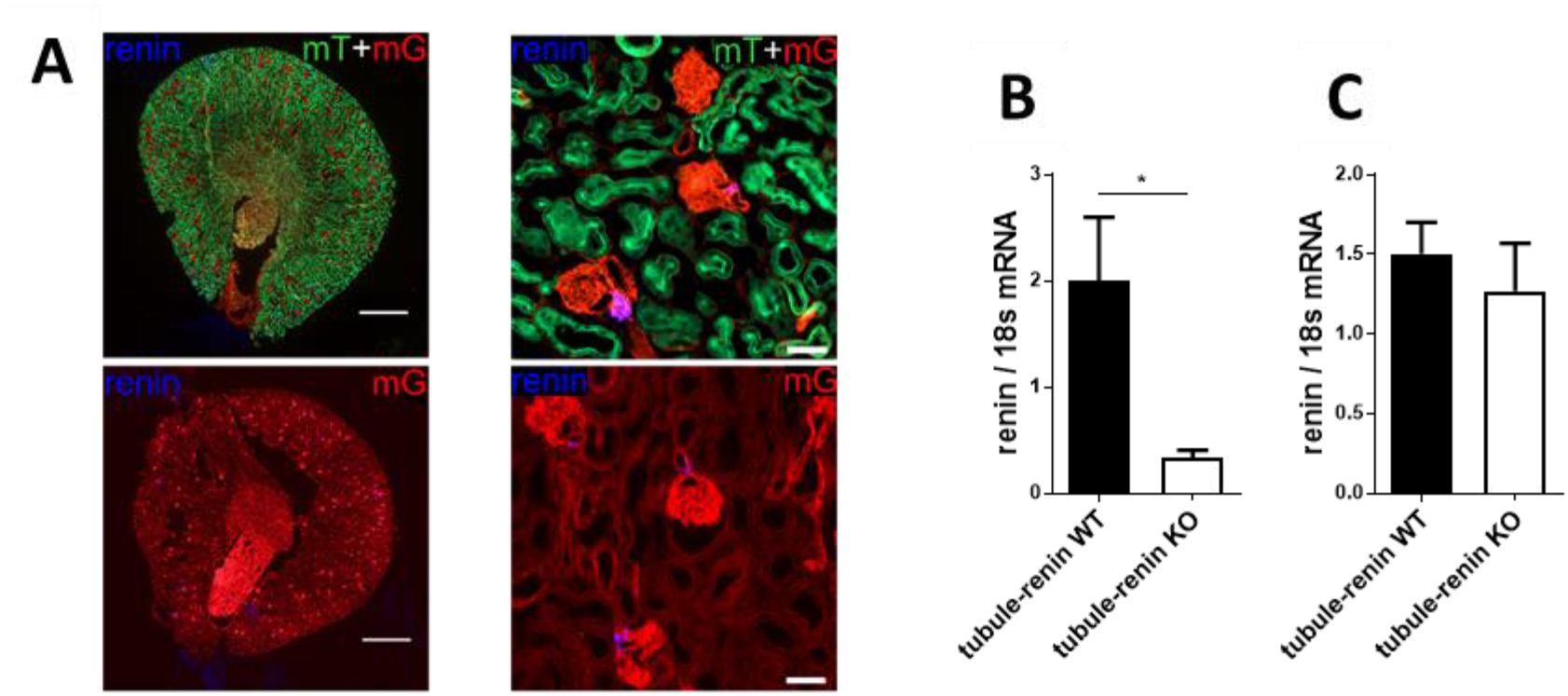
Characterization of tubule-specific renin knockout mice. (A) Pax8-TetO-Cre deleter mice were crossed with mT/mG reporter mice. Without doxycycline-induced activation of Cre-recombinase all cells show red fluorescence (lower panel). In response to doxycycline, all cells with Cre-induced recombination display green GFP-fluorescence instead of red. Costaining with a (pro)renin antibody (blue) indicate that renin expressing juxtaglomerular cells did not undergo recombination, since they show purple staining as a combination of blue (renin staining) and red (fluorescence of unrecombined reporter). (B) Renin mRNA expression in microdissected collecting ducts of tubule-specific renin knockout mice and wildtypes. (C) Renin mRNA expression in isolated glomeruli of tubule-specific renin knockout mice and wildtypes. * p < 0.05. Scale bars in A: overview: 1 mm, higher magnification: 50 μm.

Deletion of renin in the tubular system did not result in phenotypic alterations under control conditions, since no differences in body weight, urine, sodium and potassium excretion, urinary pH and arterial blood pressure were observed between tubule-renin KO and control mice (table 1).

**Table 1:** Baseline characteristics of tubule-specific renin knockout mice.

| parameter | sex | tubule-renin WT | tubule-renin KO |
| --- | --- | --- | --- |
| Body weight [g] | female | 24,9 ± 0,1 (n=14) | 25,5 ± 0,7 (n=14) |
|  | male | 33 ± 0,7 (n=16) | 32,8 ± 0,8 (n=16) |
| Weight both kidneys [mg] | female | 311 ± 11,9 (n=6) | 301,7 ± 6,1 (n=6) |
|  | male | 416,5 ± 19,4 (n=13) | 409,9 ± 16,3 (n=10) |
| Systolic blood pressure [mm Hg] | female | n.d. | n.d |
|  | male | 115,9 ± 2,7 (n=6) | 119,1 ± 4,8 (n=6) |
| Urine volume [ml / day] | female | 1,46 ± 0,13 (n=4) | 1,74 ± 0,24 (n=4) |
|  | male | 1,58 ± 0,12 (n=9) | 1,68 ± 0,20 (n=8) |
| Na<sup>+</sup>-excretion [μmol / day x g body wt] | female | 3,52 ± 0,29 (n=6) | 4,17 ± 0,36 (n=5) |
|  | male | 3,33 ± 0,36 (n=11) | 3,18 ± 0,42 (n=10) |
| K<sup>+</sup>-excretion [μmol / day x g body wt] | female | 8,96 ± 0,65 (n=6) | 8,99 ± 1,08 (n=5) |
|  | male | 10,1 ± 0,9 (n=11) | 10,86 ± 0,65 (n=10) |
| pH-urine | female | 6,1 ± 0,13 (n=4) | 6 ± 0,13 (n=4) |
|  | male | 6,4 ± 0,23 (n=8) | 6,2 ± 0,27 (n=8) |
| Plasma renin [ng angI / h per ml] | female | 58,6 ± 6,3 (n=10) | 53,8 ± 5,9 (n=8) |
|  | male | 59,8 ± 2,8 (n=15) | 57,3 ± 4,4 (n=16) |
| Hematocrit [%] | female | 49,2 ± 0,3 (n=14) | 49,1 ± 0,3 (n=14) |
|  | male | 50,1 ± 0,5 (n=16) | 49,6 ± 0,4 (n=16) |

### Marked alterations in the circulating and local renin-systems in diabetic tubule-renin KO mice

Treatment of the mice with STZ resulted in a fivefold increase in fasting blood glucose levels, marked diuresis and glucosuria in both genotypes (Fig. 3).

**Figure 3:**
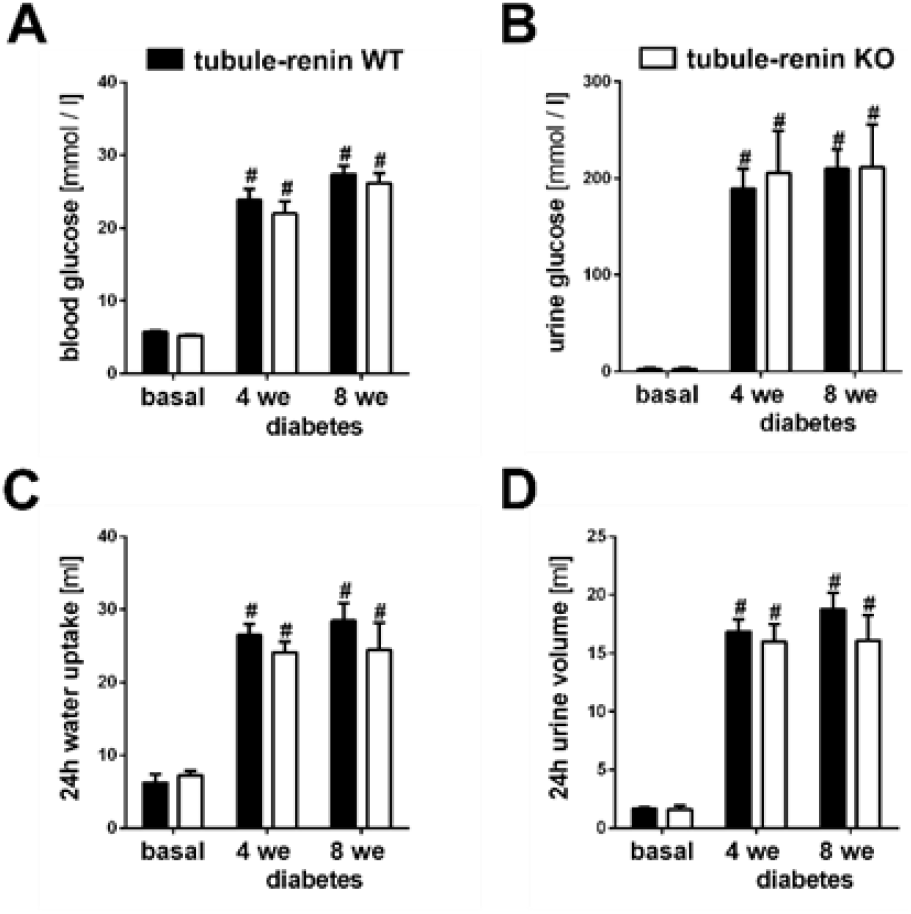
Streptozotocin induces hyperglycemia in tubule-specific renin knockout and wildtype mice. (A) Blood glucose concentration before (basal) and 4 and 8 weeks after streptozotocin (diabetes). (B) Urine glucose concentration. (C) Daily water intake. (D) Daily urine excretion. ## p < 0.001 vs. basal.

(Pro)renin immunostaining in JG-cells remained unaltered even up to 12 weeks after induction of diabetes in control and tubule-renin KO mice (Fig. 4A). (Pro)renin expression was observed in connecting tubules and collecting ducts of diabetic mice. However, while 57 % of collecting ducts of diabetic control mice were clearly (pro)renin positive, only 20 % of collecting ducts of diabetic tubule-renin KO mice showed a faint (pro)renin staining (Fig. 4B). Whether the remaining (pro)renin in tubule-renin KO mice results from incomplete deletion of the renin gene or from (pro)renin uptake cannot been discriminated from our data. Renin mRNA, active renin and prorenin tissue content had a clear cortico-medullary gradient, with high activity and concentration in the cortex and a low concentration in the inner medulla (Fig. 4C, D, E). Due to this steep gradient, contamination of samples taken from the outer medulla with tissue from the cortex will markedly affect the results. Therefore, only samples from cortex and inner medulla were analyzed. Glomerular preprorenin mRNA expression was not different between normoglycemic and diabetic mice or between genotypes (Fig 4C). In contrast, mRNA abundance was significantly stimulated by diabetes in cortical tubuli and the inner medulla from control mice, further corroborating tubular renin synthesis (Fig. 4C). This stimulation by diabetes was absent in tubule-renin KO mice, resulting in significantly lower tubular renin mRNA levels (Fig. 4C). The differences in tubular renin mRNA abundances were paralleled by alterations of the tissue prorenin concentration in cortex and inner medulla (Fig. 4E). While prorenin was significantly elevated in diabetic control mice compared with non diabetic controls, this response was significantly attenuated in the cortex and absent in the inner medulla of tubule-renin KO mice (Fig. 4E). No stimulation of tissue renin activity by diabetes occurred in any of the genotypes (Fig. 4D).

**Figure 4:**
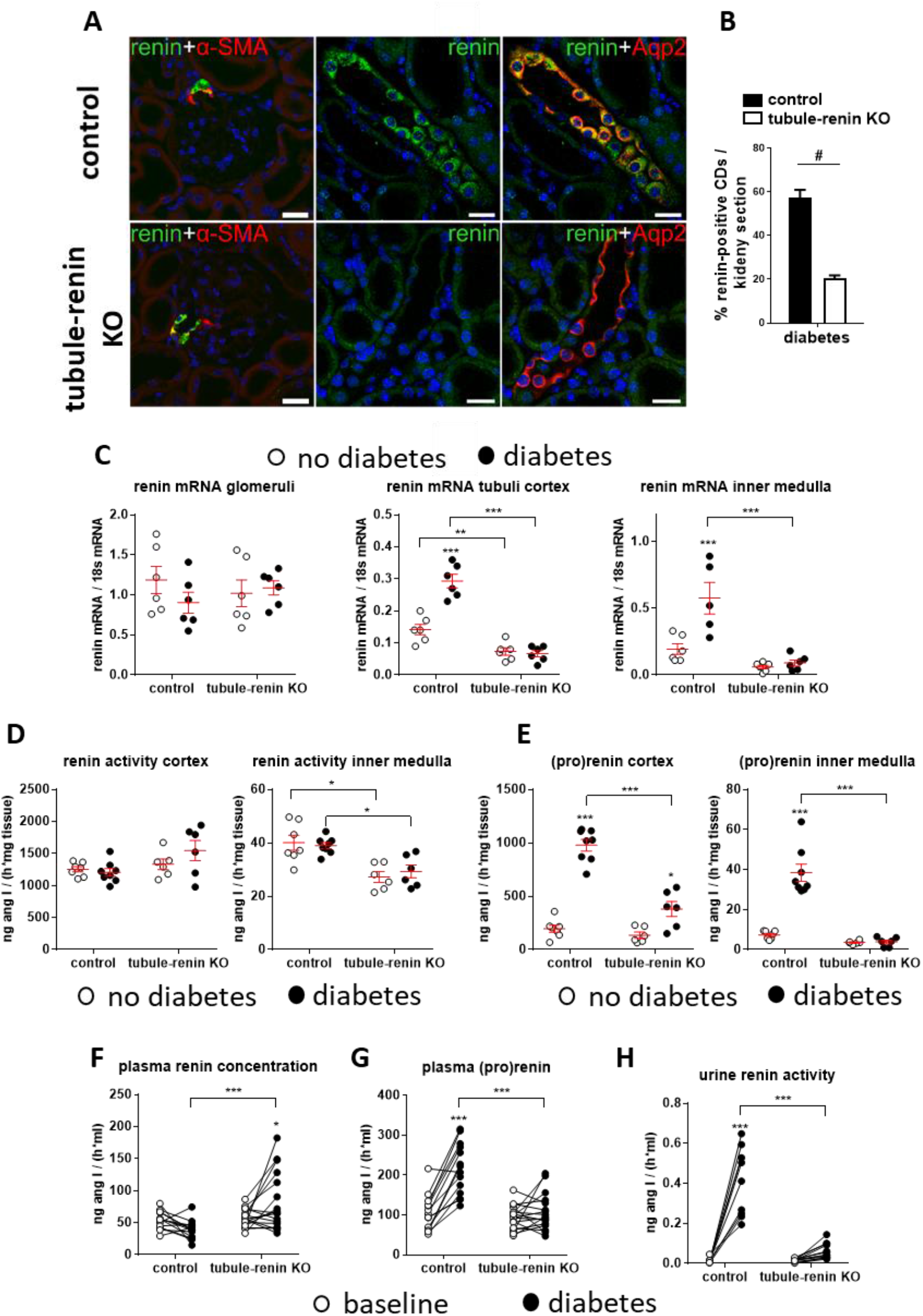
Effects of streptozotocin diabetes on parameters of the renin system. (A) (Pro)renin staining in kidneys from diabetic control mice (tubule-renin WT) or tubule-renin KO. (B) Number of (pro)renin positive collecting ducts in diabetic control and tubule-renin KO mice (n=4 each). (C) Preprorenin mRNA in non-diabetic (white circles) or diabetic (black circles) control and tubule-renin KO mice. Determination was performed in glomeruli, cortical tubules and tissue from the inner medulla. (D) Tissue renin activity in renal cortex and inner medulla. (E) Tissue prorenin concentration in renal cortex and inner medulla. (F) Plasma renin concentration before STZ (baseline) and under diabetic conditions. (G) Plasma prorenin concentration at baseline and under diabetic conditions. (H) Renin activity in the urine at baseline and under diabetes. * p < 0.05; ** p < 0.01; *** p < 0.001

Plasma prorenin concentration increased twofold in diabetic control mice compared with baseline values before STZ-treatment (104.2+12.1 ng angI/(h*ml) baseline, 215.7+17.2 ng angI/(h*ml) diabetes, Fig. 4G). This increase was completely prevented in tubule-renin KO mice (92.1+7.7 ng angI/(h*ml) baseline, 107.7+11.0 ng angI/(h*ml) diabetes, Fig. 4G), suggesting that the well-known high prorenin levels, which also occur in patients and are associated with diabetic microvascular complications (Luetscher et al., 1985; Wilson and Luetscher, 1990; Chiarelli et al., 2001), derive from renal tubules. Also in line with results from diabetic patients (Fernandez-Cruz et al., 1981; Price et al., 1999; Chiarelli et al., 2001), plasma renin concentration tended to decrease in diabetes in control mice (Fig. 4F). Unexpectedly, plasma renin concentration was significantly stimulated increased in tubule-renin KO mice (Fig. 4F). Renin activity in the urine was barely detectable at baseline before STZ application but markedly increased under diabetic conditions (Fig. 4H). This effect was blunted in tubule-renin KO mice, indicating renin release from the tubules into the urine (Fig. 4H). No prorenin was detectable in the urine in any of the groups (not shown).

### Tubule-specific renin KO mice have reduced albuminuria and glomerular mesangial expansion

Diabetes mellitus was accompanied by enhanced albumin excretion in tubule-renin control and KO mice. However, albuminuria was significantly lower in diabetic tubule-renin KO mice compared with controls (control mice: baseline 40.1+2.8, diabetes 173.2+16.6 μg/mg creatinine, tubule-renin KO mice: baseline 36.6+3.0, diabetes 113.7 + 9.9 μg/mg creatinine, Fig. 5A). Microalbuminuria might result from either enhanced glomerular filtration or from reduced tubular reabsorption. Diabetic tubule-renin KO mice had a significantly lower albumin staining in proximal tubules compared with the tubule-renin control mice, indicating that the amelioration of albuminuria in diabetic KO mice did not result from stimulated tubular albumin reabsorption but from reduced glomerular albumin filtration (Fig. 5B, 5C). STZ treatment induced glomerular hypertrophy to a similar extend in both genotypes (Fig. 5D, 5E), whereas mesangial expansion was only detectable in tubule-renin control mice, but not in their KO littermates (fig. 5F). These differences in glomerular histology did not result in parallel changes in glomerular filtration rate. As shown in figure 5G, GFR tended to be higher in tubule renin control mice compared with knockouts, however this numerical difference did not reach statistical significance.

**Figure 5:**
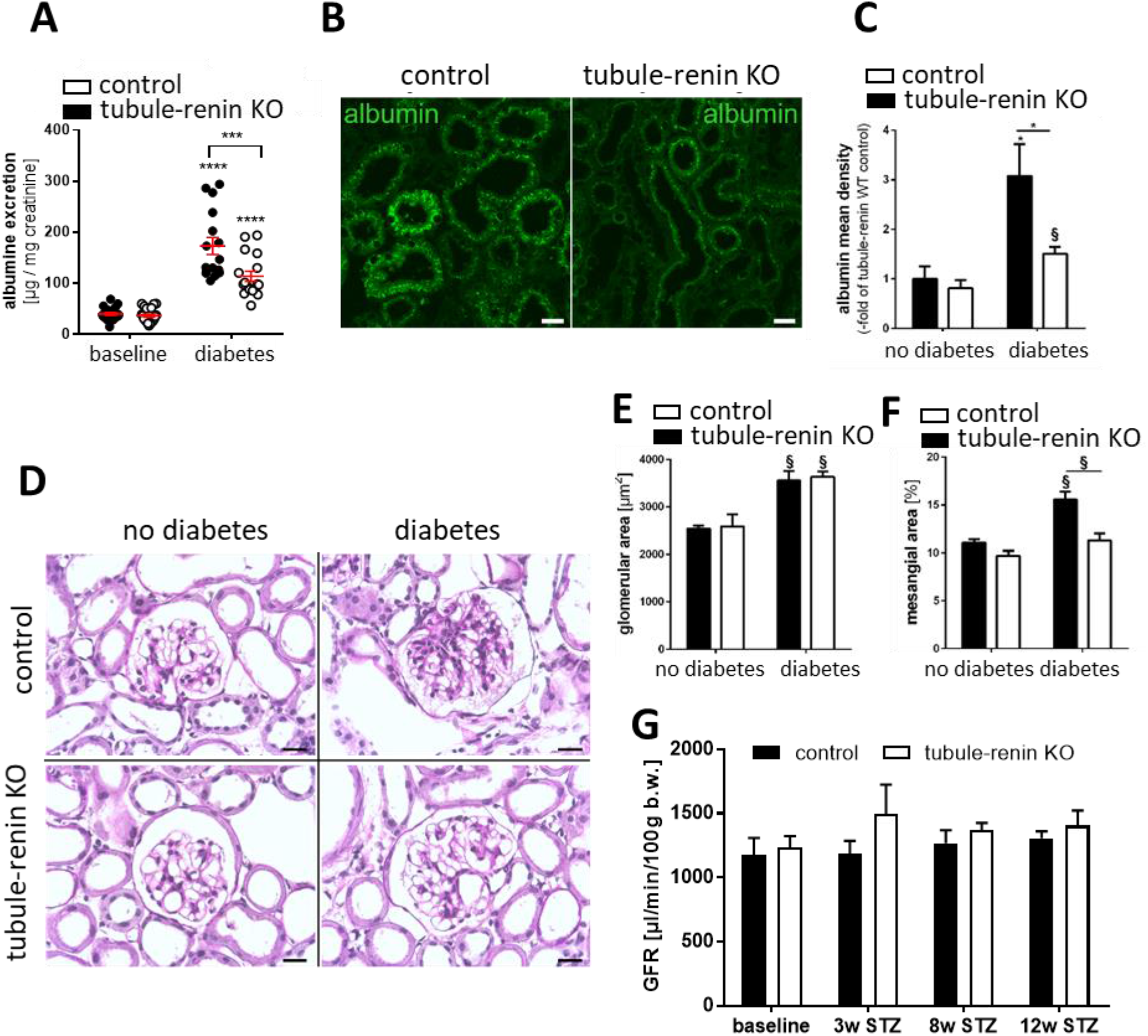
Effects of STZ-diabetes on albuminuria, glomerular hypertrophy and glomerular filtration rate. (A) Albuminuria in control mice (tubule-renin WT) or tubule-renin KO under normoglycemic baseline conditions or STZ diabetes. (B) Albumin staining in kidneys from diabetic control or tubule-renin KO mice. (C) Determination of albumin fluorescence density (n=6 each). (D) Histological examination of glomeruli. (E) Stereological assessment of glomerular area and mesangial area (F, as percent of glomerular area). (G) Glomerular filtration rate at baseline and 3, 8, and 12 weeks after induction of diabetes with streptozotocin (STZ). * p < 0.05; §

### Diabetic interstitial fibrosis, inflammation and capillary rarefaction are ameliorated in tubule-renin KO mice

Masson-Goldner staining showed expansion of the fibrotic area in diabetic kidneys of both genotypes (Fig. 6A). Unbiased, automatic quantification revealed that the fibrotic response was moderately, but significantly attenuated in tubule-renin KO mice (Fig. 6B). Similarly, the number of intrarenal macrophages (stained with F4/80 as marker) significantly increased in diabetic kidneys and, again, this response was attenuated in mice with deletion of renin in the tubular system (Fig. 6C, D). Finally, staining of endothelial cells (antibody directed against CD31) and quantification of positively stained capillaries showed a decrease in capillary density in diabetic kidneys of wildtype mice. This effect was completely abolished in tubule-renin KO mice (Fig. 6E, F).

**Figure 6:**
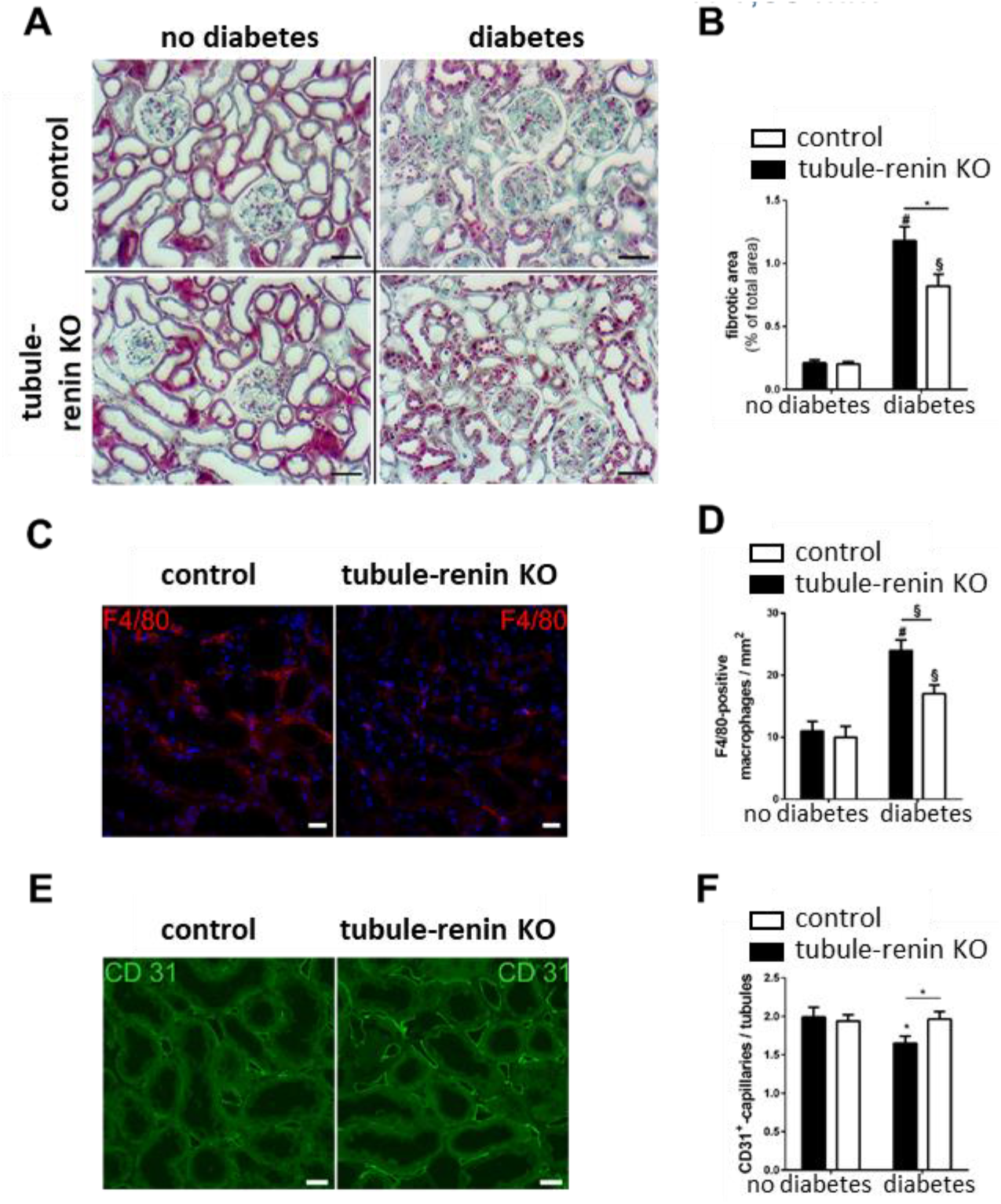
Effects of STZ-diabetes on renal fibrosis, inflammation and capillary rarefaction. (A) Masson-Goldner staining in kidneys from control mice (tubule-renin WT) or tubule-renin KO under normoglycemic or diabetic conditions. Collagen deposits show green/blue staining. Scale bar: 50 μm. (B) Quantification of fibrotic area. (C) Staining of macrophages (marker protein F4/80) in kidneys from diabetic control or tubule-renin KO mice. Scale bar: 20μm. (D) Quantification of macrophage number per area. (E) Staining of endothelial cells (marker protein CD31) in peritubular capillaries. Scale bar: 20μm. (F) Number of peritubular capillaries in kidney section from control mice or tubule-renin KO under normoglycemic or diabetic conditions. For normalization, number of capillaries were related to the number of tubules in the respective kidney section. * p < 0.05; § p < 0.01; # p < 0.001.

### Glucose stimulates renin synthesis in mouse collecting duct cells in a concentration dependent manner

To test whether glucose might directly stimulate renin expression in collecting duct cells, we used the M-1 cell line. Cells were either cultured with a medium containing 5 mM glucose (control) or with a medium to which further glucose was added so that the final concentration of glucose was 10, 20 or 30 mM, hereby mimicking the blood glucose concentration in STZ diabetes (see Fig. 3). In fact, abundance of preprorenin mRNA was stimulated in a concentration dependent manner by glucose (Fig. 7A). Stimulation was not observed in response to application of mannitol, indicating that the effects of glucose on renin expression are not mediated by changes in osmolarity (Fig. 7B). M-1 cells did not only synthesize but also secrete renin, indicated by the fact that renin activity and prorenin in the supernatant were elevated in response to glucose (Fig.7C, D).

**Figure 7:**
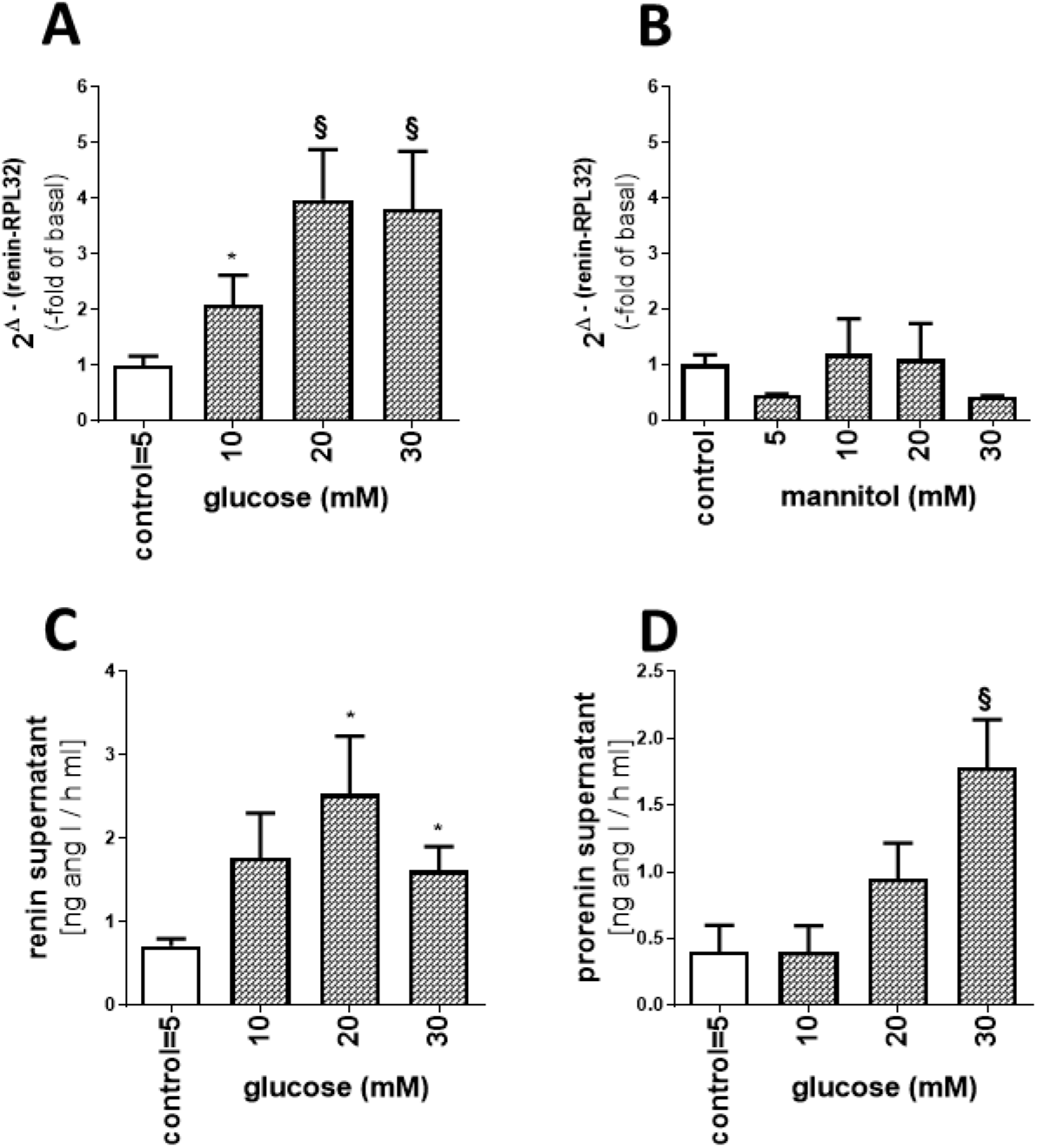
Effects of increasing glucose concentrations on renin synthesis in collecting duct cells. (A) Stimulation of renin mRNA expression by glucose. For normalization, renin mRNA abundance was related to mRNA concentration of RPL32. (B) Elevation of osmolarity by the addition of mannitol did not stimulate renin expression. (C) The concentration of active renin was significantly stimulated by glucose. (D) Stimulation of prorenin concentration in the supernatant by high glucose concentration. * p < 0.05; § p < 0.01.

## Discussion

As one main result, the present study provides evidence that distal tubules and collecting ducts are sources of prorenin and markedly contribute to elevated concentrations of prorenin in renal tissue and plasma in diabetes. This conclusion is based on several lines of evidence: I. Collecting ducts and distal tubules show clear (pro)renin staining in kidney from diabetic mice and patients. This result is in agreement with previous studies in rats and mice (Kang et al., 2008; Tang et al., 2019). However, while the first study concluded that tubular renin protein results from tubular synthesis, the latter concluded that it derives from tubular uptake. Although direct evidence for tubular uptake of renin in collecting ducts is not presented in the latter paper, data from urinary renin excretion in diabetic patients and very elegant and sophisticated pharmacological experiments in diabetic mice suggest that renin, which enters proximal tubules by glomerular filtration, is reabsorbed in proximal tubules (Tang et al., 2019).

Since megalin expression in proximal tubules is reduced in diabetic kidneys, defective renin reabsorption could result in high urinary renin levels in distal nephron segments and reabsorption of renin in collecting ducts (Tang et al., 2019). Renin and prorenin are smaller than albumin, can be filtered in the glomeruli and their glomerular sieving coefficient increases in glomerular diseases (Roksnoer et al., 2016). Moreover, it had been shown previously that patients with defective or mice with pharmacologically blocked proximal protein reabsorption have high renin concentrations in the urine (Mazanti et al., 1988; Roksnoer et al., 2016). Tang et al. extended these studies to diabetic mice and blocked proximal protein uptake by infusion of lysine, resulting in a marked increase in urinary renin excretion (Tang et al., 2019). Unfortunately, in none of those studies renin staining was performed in order to investigate whether blockade of renin uptake in proximal tubules in fact induces enhanced renin staining in collecting ducts. Thus, clear direct evidence for tubular uptake of renin in collecting ducts is missing to date. The data of our study cannot exclude that (pro)renin staining in collecting ducts at least partially derives from renin uptake. (Pro)renin staining was markedly reduced, but not completely abolished in collecting ducts of tubule-renin KO mice. This residual (pro)renin protein might either result from incomplete deletion of renin, as indicated by the residual renin mRNA in collecting ducts from tubule-renin KO mice, or from renin uptake from the tubular fluid. However, the fact that tubular (pro)renin staining was reduced in tubule-renin KO mice in which plasma renin activity was markedly elevated, what in turn should result in higher renin levels in the tubular fluid, argues against major contribution of tubular uptake to renin staining in collecting ducts. II. Direct demonstration of tubular renin mRNA in diabetic kidneys by in situ hybridization in our and a previous study (Lai et al., 1998) and upregulation of renin mRNA in isolated cortical tubuli and inner medullary tissue underline tubular renin mRNA expression and local formation of renin. It has to be mentioned that Tang et al. did not find upregulation of renin mRNA in microdissected collecting ducts, what again argues against local formation. This discrepancy between our mRNA data and the data from Tang et al. cannot be explained at the moment, but might result from the different isolation procedures of tubuli or the different mouse strains or sexes (Tang et al., 2019). III. The most convincing evidence for tubular production of prorenin comes from our *in vivo* study in tubule-specific renin knockout mice. Reduction of tubular (pro)renin protein and mRNA expression, blunted increases in tissue prorenin concentration and plasma prorenin concentrations clearly argue for synthesis and release of prorenin from tubules of diabetic mice and cannot be explained by altered tubular uptake.

Tubule-renin KO mice had no renal phenotype at baseline, suggesting that tubular renin does not play a functional role under control conditions. In line with this assumption, (pro)renin staining was not observed in non diabetic mice, most likely due to (pro)renin levels below the detection limit of our immunofluorescence approach. Alternatively, the deficiency of tubular renin could be compensated by renin from JG cells. Plasma renin activity was not different between tubule-renin KO and controls under control conditions. However, while plasma renin activity was not significantly altered in diabetes compared to non-diabetic baseline conditions in control mice, it was markedly stimulated by diabetes in tubule-renin KO mice. This unexpected result might be explained by a compensatory increase in active renin as a result of low prorenin concentrations in renal tissue and plasma. For instance and as discussed below in more detail, binding of prorenin to the (pro)renin receptor (PRR) in collecting ducts, stimulation of NaCl reabsorption and elevation of blood pressure can be induced (Ramkumar and Kohan, 2019). Accordingly, reduction of local or systemic prorenin in tubule-renin KO mice could result in salt deficiency and low blood pressure, especially in the situation of markedly enhanced diuresis as in our STZ model. This threatening situation may be counterbalanced by stimulation of plasma renin activity, which would lead to stimulation of NaCl reabsorption and stabilization of blood pressure via increased angiotensin II concentrations and subsequent aldosterone release.

Another main result of our study is the marked protection from diabetic kidney damage in tubule-renin KO mice. In general, angiotensin II is considered as disease promoting factor in diabetic nephropathy and, accordingly, AT1 receptor antagonists have been proven to be effective in slowing down the progression of the disease. In light of these well known facts, renoprotection in tubule-renin KO mice is surprising, since these mice have markedly elevated plasma renin activity, what most likely results in high angiotensin II concentrations. Therefore, the amelioration of diabetic renal damage is presumably attributable to low prorenin levels in diabetic tubule-renin KO mice, what further underlines an important pathophysiological role of prorenin in disease progression as has been suggested in early studies (Luetscher et al., 1985; Wilson and Luetscher, 1990). Prorenin, and also renin, can bind to the (pro)renin receptor (PRR) that is expressed in many tissues including the kidney. Binding of prorenin to the PRR induces a conformational change of prorenin resulting in activation of prorenin without the cleavage of the prosegment (Nguyen et al., 2002; Nguyen, 2007). Therefore, binding of prorenin to the PRR is accompanied by local production of angiotensin II. In addition to its role in angiotensin II formation, binding of prorenin to the PRR activates intracellular signaling pathways such as ERK1 and 2 (extracellular-signal-regulated kinase 1 and 2) and MAPK (p38 mitogen-activated protein kinase), hereby inducing profibrotic mechanisms (Nguyen et al., 2002; Nguyen, 2007). Finally, the PRR is an integral component of the vacuolar H^+^-ATPase that is expressed in all cell types and contributes to acidification of organelles (Forgac, 2007). Accordingly, genetic deletion of PRR in the tubular system affected urinary acidification (Trepiccione et al., 2016). Moreover, several studies showed that PRR is expressed in renal collecting ducts and significantly contribute to the urinary concentrating abilities of the kidney, to sodium balance and blood pressure control (comprehensively reviewed in (Arthur et al., 2021)). As mentioned above, low prorenin concentrations observed in tubule-renin KO mice might therefore result in an activation of the circulating active renin in order to compensate for the missing tubular effects of prorenin. PRR is also expressed in podocytes and mesangial cells and upregulated by diabetes and high glucose concentrations. Silencing of PRR by specific siRNA ameliorated podocyte integrity and function in hyperglycemic conditions (Siragy and Huang, 2008; Li and Siragy, 2014). Consistent with a disease-promoting effect of the PRR, pharmacological blockade of PRR ameliorated renal injury in diabetic rats (Matavelli et al., 2010). Therefore, it seems very plausible that in our study, low prorenin concentrations in diabetic tubule-renin KO mice resulted in the observed reduced diabetic renal injury.

In conclusion, our study identified renal tubules as source for tissue and plasma prorenin in diabetes. Moreover, it provides evidence that tubular prorenin is a major disease aggravating factor in diabetic kidney damage and, therefore, allows a better understanding of the pathogenesis of diabetic nephropathy.

